# Structural and functional brain asymmetry in relation to heterogeneous causes of *situs inversus totalis*

**DOI:** 10.1101/2025.11.04.686509

**Authors:** Meng-Yun Wang, Nicole Ng, Else Eising, Simon E. Fisher, Guy Vingerhoets, Clyde Francks

**Affiliations:** Language & Genetics Department, Max Planck Institute for Psycholinguistics, Nijmegen, The Netherlands; Donders Institute for Brain, Cognition and Behaviour, Radboud University, Nijmegen, The Netherlands; Department of Experimental Clinical and Health Psychology, Ghent University, Ghent, Belgium; Department of Medical Neuroscience, Radboud University Medical Center, Nijmegen, The Netherlands

**Keywords:** Situs inversus, functional brain asymmetry, structural brain asymmetry, laterality, genome sequencing, brain imaging, cilia

## Abstract

Various aspects of brain organization differ between the left and right hemispheres. Clues to the developmental origins of these asymmetries may be gained through associations with *situs inversus totalis* (SIT), a rare condition in which visceral organs are reversed on the left-right axis. In the largest previous brain imaging analyses of SIT (38 cases, 38 controls from Belgium), typical functional asymmetries such as left-hemisphere language dominance were mostly unaltered, but various aspects of asymmetrical cerebral structure - *petalia*, bending, and posterior venous anatomy – were often reversed in this condition. SIT can be a monogenic trait that arises from rare genetic variants, usually affecting motile cilia which help to create the embryonic left-right body axis. However, most SIT cases do not have obvious genetic causes and may arise from environmental or random effects during embryogenesis. We sequenced the genomes of 23 SIT cases and 23 controls from the Belgian brain imaging dataset and pooled with prior data from 15 cases and 15 controls. We aimed to discover whether there are altered brain asymmetries in SIT cases with disruptive DNA variants in ciliary genes, or in other types of genes, as compared to genetically unsolved cases. In total, 19 cases had likely causal variants affecting ciliary function, while 19 cases remained genetically unsolved. Functional and structural brain asymmetries were not significantly different in genetically solved versus unsolved SIT cases. Therefore, functional brain asymmetries seem largely independent of known mechanisms of visceral *situs* formation, while structural brain torque is altered in SIT regardless of the presence or absence of overt genetic causes.

## Introduction

The human brain is asymmetrical both functionally and structurally ^1-3^. Prominent examples of functional asymmetry include left hemisphere dominance for language processing and skilled manual actions in most people ^4,5^, and relatively greater right hemisphere engagement for spatial attention and face recognition ^6,7^. Functional asymmetries are likely to have deep evolutionary roots, adapted from ancient neurophysiological and behavioural asymmetries in our vertebrate ancestors ^8^.

In terms of structure, there is a cerebral skewing or ‘torque’ ^9,10^, which manifests as *petalia* and bending ^11-13^. *Petalia* refer to asymmetrical protrusions of the cerebral hemispheres along the antero-posterior axis, typically manifesting as right-frontal and left-occipital extensions. Bending involves a displacement of brain tissue across the midline – where typically the left occipital lobe intrudes into the contralateral hemisphere’s space ^11-13^. Other structural brain asymmetries include region-specific hemispheric differences of cerebral cortical surface area and/or thickness ^14,15^, some of which relate to torque ^16,17^, as well as asymmetries of subcortical structural volumes ^18^ and white matter tracts ^19^. In addition, asymmetry is present in the posterior venous system, where the right transverse sinus receives flow from the superior sagittal sinus, while the left transverse sinus receives flow from the smaller, straight sinus, resulting in a marked rightward asymmetry of transverse sinus volume in most people ^12,13^.

While there are typical forms of functional and structural brain asymmetries in the general population, there is also considerable variation. For example, 5–15% of people show bilateral or right-hemisphere dominance for speech production ^20,21^, and roughly 30% of people show an absent or mirror-reversed torque pattern compared to the majority form ^17^. Different aspects of brain structural and/or functional asymmetry can vary largely independently of each other, such that brain asymmetry may be considered a complex, multivariate phenomenon with components of variation that are dissociable ^9,22,23^. Reduced or absent brain asymmetries are associated with lower cognitive performance and/or neuropsychiatric disorders ^24-26^, which is consistent with a role for hemispheric specialization in optimal brain functioning.

Precursors of adult human brain asymmetries and handedness are already observable as early as the first and second trimesters of pregnancy ^27-30^, indicating that the brain’s left-right axis forms during embryonic patterning. Recently, population variation in brain and behavioural asymmetries has been associated with numerous genetic variants through large-scale genome-wide association studies ^31-33^. The implicated genes notably include several that encode microtubule components or associated proteins ^34-36^. However, the mechanisms by which these genes affect brain asymmetries, or the extent to which they are involved in patterning the embryonic left-right brain axis, remain uncertain (see also the Discussion).

Further possible clues to the developmental bases of brain asymmetries may come from their associations with visceral organ situs ^7,13^. The internal organs in the thorax and abdomen are typically positioned according to an asymmetrical layout known as *situs solitus* (SO). For example, the stomach and spleen are positioned toward the left side of the body, with the liver toward the right ^37^. For visceral organ asymmetries, core developmental mechanisms are better understood than for the brain: The unidirectional movement of motile cilia in the node, a transient structure of early mammalian embryos, creates a leftward flow of fluid, likely triggering an asymmetry of calcium ion flow into cells around the node, driving differences of gene expression on the two sides, which ultimately specify typical situs of the visceral organs ^38^.

Nonetheless, rare forms of situs abnormalities are found in the population, which can arise from loss-of-function variants in more than 100 different genes involved in left-right axis formation ^37^. When all thoracic and abdominal organs are reversed on the left-right axis, the condition is called *situs inversus totalis* (SIT). The prevalence of SIT is estimated at around 1/10000 ^37^. Rare DNA variants that disrupt motile cilia are prominent causes of SIT ^39^, and those cases often co-occur with primary ciliary dyskinesia (PCD), where impaired movement of cilia causes deficits in mucus clearance from the lungs and can lead to respiratory infections ^40,41^. However, in more than half of SIT cases no genetic basis has been identified ^40^, and environmental factors may play a role, such as maternal smoking or other pregnancy risks ^37^. The possibility of random effects in very early embryonic patterning must also be considered ^42^. This might involve, for example, chance fluctuations in the concentrations of key patterning molecules on the left and right sides, while developmental trajectories are being set ^43^.

Intriguingly, visceral organ reversal in SIT does not usually coincide with a mirror reversal of functional brain lateralization ^6,7,12^. For example, language lateralization in individuals with SIT generally follows the typical pattern of left-hemisphere dominance ^6,7,12,44,45^. Furthermore, left-handedness was not found to be more common in individuals with SIT and PCD than in unaffected controls ^46^.

Nonetheless, a recent study from Belgium with 38 SIT individuals – the largest dataset of SIT with brain imaging - observed that unusual patterns of hemispheric crowding of functions and mirror-reversal of functional organization were more common in SIT (48%) than in controls (30%) ^7^. Two typically left-lateralized functions (speech production and manual praxis) and two typically right-lateralized functions (spatial attention and face recognition) were evaluated with functional Magnetic Resonance Imaging (fMRI). The case-control group difference did not achieve statistical significance, but the possibility was raised that individual cases may have disrupted developmental mechanisms that affect both visceral and functional brain asymmetries. This is plausible given that SIT can arise from loss-of-function variants in numerous different genes, as well as from non-genetic causes. In addition, left-handedness was found to be significantly more common in the Belgian SIT cases (26%) than in the general population (10.6%) ^7^, again suggesting that an etiologic subgroup of SIT individuals might have disrupted functional brain laterality.

In terms of structural brain asymmetry, the evidence for left-right reversals accompanying SIT is now clear ^12,13,44,45,47,48^. The largest study was again of the Belgian dataset (38 SIT, 38 controls), where significant group average reversals were found for occipital and frontal petalia and occipital bending, as well as transverse sinus asymmetry ^13^. Therefore, there appears to be a developmental link between these aspects of structural brain asymmetry and visceral organ asymmetry, but one which is more tenuously linked to functional brain asymmetry.

In an earlier subset of the Belgian dataset (15 SIT cases, 15 controls), genome sequencing was performed which resulted in likely causative genetic variants being identified in 10 of the cases ^49^. Eight of these cases carried disruptive variants in genes known to cause PCD and SIT, while two cases had variants in genes known to cause SIT without PCD, but still in cilia-related genes (*PKD1L1, CFAP52*). Five of the SIT cases had no obvious monogenic basis for their condition, including three left-handers, such that a genetic link of SIT to behavioural asymmetry could not be established. Brain imaging data, however, were not considered in that study ^49^.

In the present study, we first sequenced the genomes of 23 more SIT cases and 23 controls from the Belgian dataset to identify likely causal DNA variants, and then pooled with the earlier subset ^49^ for a total of 38 SIT cases and 38 controls. We aimed to investigate whether the causal heterogeneity of SIT relates to altered functional and structural brain asymmetry, by integrating genetic and brain imaging data for the first time in this dataset.

## Methods

### Participants

This study included 38 participants with SIT and 38 matched controls, who were recruited in two stages. In stage I, 15 SIT participants were recruited from Ghent University Hospital and Middelheim Hospital Antwerp, through radiological database searches ^6,12,49^. Ethical approval was obtained from the medical ethical committee of Ghent University Hospital (project number 2012/073). In stage II, 23 SIT participants were recruited via radiological protocol screening across six Flemish general hospitals ^7,13^. Ethics approval was again obtained from the medical ethical committee of Ghent University Hospital (Belgian registration number B670201837416) as well as from the local medical ethical committees of the participating hospitals before recruitment was started. In each stage, control individuals equal to the number of cases were also recruited via social networks and word of mouth, and were further matched for sex, age, handedness, and education (Table 1). All participants gave informed consent for DNA sample collection, genomic analysis, and brain MRI to investigate body and brain asymmetries.

**Table 1.** Summary information of the participants. SIT: *situs inversus totalis*. PCD: primary ciliary dyskinesia. Categorical variables are given as counts and percentages. Continuous variables are given as means and standard deviations.

|  | Total |  | Stage I |  | Stage II |  |
| --- | --- | --- | --- | --- | --- | --- |
|  | SIT (38) | Controls (38) | SIT (15) | Controls (15) | SIT (23) | Controls (23) |
| Female (%) | 22 (58%) | 22 (58%) | 7 (47%) | 7 (47%) | 15 (65%) | 15 (65%) |
| Age (years) | 41.05 ± 17.04 | 40.92 ± 16.15 | 33 ± 10.1 | 33 ± 10.0 | 42.1 ± 16.1 | 42.3 ± 15.9 |
| Education (years) | 12.05 ± 2.10 | 12.16 ± 2.03 | 12.9 ± 2.3 | 12.9 ± 1.6 | 12.4 ± 2 | 12.8 ± 2 |
| PCD diagnosis | 8 (21%) | 0 | 6 (40%) | 0 | 2 (9%) | 0 |
| Right-handed | 26 (68%) | 30 (79%) | 9(60%) | 11 (73%) | 17 (74%) | 19 (83%) |

In both stages, SIT status was confirmed through radiological or MRI evidence of complete visceral reversal. Brain MRI scanning was performed at Ghent University Hospital. Among the 38 SIT participants from both stages combined, eight were identified as likely cases of PCD or Kartagener syndrome, which refers to the clinical triad of *situs inversus*, chronic sinusitis, and bronchiectasis (permanent enlargement of the airways) ^50,51^, based on clinical records and PICADAR scores ^6,7,12,52^. Three SIT participants from stage I had records of congenital heart disease requiring surgical intervention, as documented in their radiological reports. This aligns with the known association between laterality defects and cardiac abnormalities ^37^.

The remaining SIT participants in stage I reported no major medical issues; they had undergone radiological examinations for diverse reasons, including gastric complaints, general fatigue, minor accidents, or unspecified reasons. In stage II, various participants—both SIT and controls—reported past neurological (e.g., migraine, essential tremor), psychiatric (e.g., depression, anorexia), or developmental conditions (e.g., ADHD, dyslexia), but these conditions were comparable in type and frequency between cases and controls, and did not interfere with behavioural assessments or imaging procedures ^7,13^. The cognitive level of individuals with SIT was comparable to the general population ^6,7^. After consideration of Edinburgh Handedness Inventory scores ^53^ together with self-reports of hand switching or inconsistent hand preference, the combined SIT sample consisted of 28 right-handed, 9 left-handed, and 1 ambidextrous individual.

Further details on inclusion and exclusion criteria, and the clinical and behavioural assessments, can be found in earlier publications on this dataset ^6,7,12^. Summary information on demographics, education, PCD and handedness can be found in Table 1.

### Genome sequencing, variant calling and filtering

Details of DNA sequencing, variant calling, annotation, and filtering for stage I individuals were described previously ^49^.

For the present study, whole genome sequencing of 23 additional SIT cases and 23 controls (stage II individuals) was performed by Novogene Co. Ltd. on the DNBseq platform (150 bp paired-end reads), on DNA extracted from saliva using Oragene kits. Reads were aligned to the human reference genome (build 38, decoy version) using Burrows-Wheeler Aligner (BWA v0.7.17)^54^, and processed with Sambamba (v1.0.0)^55^, Picard (v2.18.9) (https://broadinstitute.github.io/picard/), and Genome Analysis Toolkit (GATK v4.3.0) ^56^ according to the GATK Best Practices. Variants were called using GATK HaplotypeCaller and filtered with the GATK VariantFiltration module based on the following criteria, which were applied separately for single nucleotide variants (SNVs) and insertion-deletions (indels). For SNVs, variants were excluded if they met any of the following thresholds: Quality by Depth (QD) < 2, Fisher Strand Bias (FS) > 60, Mapping Quality (MQ) < 40, HaplotypeScore > 13, Mapping Quality Rank Sum Test (MQRankSum) < −12.5, or Read Position Rank Sum Test (ReadPosRankSum) < −8. For indels, variants were excluded if QD < 2, FS > 200, or ReadPosRankSum < −20. This yielded a combined total of 266,596,901 retained SNVs and indels (mean 5,795,585 variants per genome).

Variants were annotated with Annovar ^57^ using several databases: ensGene 45 (FASTA sequences for all annotated transcripts in Gencode v45 Basic collection; last update was 2024-01-11 at UCSC), dbSNP 151 ^58^, dbnsfp (version 4.7a) ^59,60^, gnomAD (version 4.1) ^61^, ClinVar (20250217) ^62^.

Following annotation, both variant-level and gene-level filtering steps were applied. At the variant level, only exonic and splicing variants were retained, while synonymous SNVs, variants labeled as “unknown,” and un-annotated variants were excluded. Variants with Combined Annotation Dependent Depletion (CADD) Phred scores >= 20 were retained as likely to be disruptive to protein function ^63^. This was based on CADD v1.7 that was updated in 2024: These scores integrate multiple functional annotations into one metric using Machine Learning-based scoring and classification of genetic variants, and can be derived for both SNVs and indels. Additionally, we retained missense SNVs with CADD < 20 when there was a pathogenic prediction by AlphaMissense ^64^; variants with benign AlphaMissense predictions were excluded. Variant quality filters were also applied: SNVs with read depth (DP) below 7 or genotype quality (GQ) below 20, and indels with DP < 10 or GQ < 20, were excluded. Heterozygous variants were excluded if the allelic balance (AB) was more extreme than 3:1 or 1:3 in terms of read depth.

### Recessive/X-linked and dominant models

As SIT and PCD are rare in the population and their monogenic causes are mostly known to be recessive ^37^, we first queried the stage II data under a recessive model (as we had previously for stage I ^49^). Variants with a population allele frequency (gnomAD_AF) greater than 0.005 were removed. Then, at the gene level, only genes that had at least two retained variants in a given SIT case, and that were not detected in phase, were retained as possible causes (this step excluded genes unless they had homozygous disruptive variants or potential compound heterozygous variants). In addition, we retained genes on chromosome X in male SIT cases when they had at least one retained variant. Genes meeting these criteria in at least one control were further excluded as possible causal genes in cases (this step removed genes that are affected by many rare disruptive variants in the population and/or affected by frequent sequencing or variant calling errors in our pipeline).

According to these criteria, stage II SIT cases had a mean number of 6.48 genes in their genomes (range 1-25) affected by biallelic or X-linked hemizygous variants. This compares to stage I SIT cases that had a mean number of 8.4 genes in their genomes (range 5-15) affected by biallelic or X-linked hemizygous variants ^49^. Note that the genome sequencing technology was different for the two stages, as well as versions of variant calling and annotation software and databases ^49^.

To identify the most likely single causal gene per SIT individual, we then compared their list of genes under the recessive/X-linked model to a list of genes that we compiled using terms to search the ClinVar database (https://ftp.ncbi.nlm.nih.gov/pub/clinvar/gene_condition_source_id) ^62^ and Mouse Genome Database 2025 ^65^: “situs inversus”, “heterotaxy | heterotaxia | situs ambiguus”, “PCD | ciliary dyskinesia | Kartagener syndrome”, “left-right”, and “asymmetry | laterality”. To this list, we also added genes from the literature focusing on situs inversus and left-right asymmetry: these included a candidate gene list compiled for our previous genetic study of stage I cases in relation to handedness ^49^, plus three recent studies of laterality disorders and/or PCD in humans ^66-68^, and large-scale genome-wide association studies of structural brain asymmetry and handedness ^31,32^.

In total, there were 351 unique genes in the candidate list (Table S1 lists the source for each gene). Finally, when matching genes were identified between our mutated gene list from SIT cases and our candidate gene list, we verified the known mode of inheritance and phenotypic associations through literature search (PubMed) and database queries for the implicated genes (OMIM ^69^, Clinvar ^62^, Genecards ^70^, Mouse Genome Database ^65^).

As performed previously for stage I ^49^, for the stage II SIT cases not solved under a recessive/X-linked model, we then considered a dominant model. Here we applied a stricter threshold for allele frequency, whereby variants with gnomAD_AF > 0.00005 were excluded. At the gene level, genes were retained as possibly causative in cases if they had at least one variant remaining (in heterozygous, homozygous or hemizygous form), except that genes meeting these criteria in more than one stage II control were further excluded as possible causes in cases, again to help exclude genes that carry many rare disruptive variants in the population, and help remove pipeline errors. This yielded a mean of 65 genes per stage II SIT case, range 31-97 under a dominant model, compared with 41.8 genes per stage I case, range 30-66. These genes were again compared to our candidate gene list, and matches were again followed up with literature and database queries (as described for the recessive model above in this section), to verify the mode of inheritance and known phenotypic associations.

### Gene set enrichment analysis

To test whether a list of genes contained functionally related genes when compiled across participants, we performed Gene Set Enrichment Analysis (GSEA) using the g:Profiler2 package (version 0.2.3) ^71^. Three databases from the genome ontology (GO) ^72^ were tested, namely biological process (BP), cellular component (CC), and molecular function (MF) ^73^. The date release of these three databases was “annotations: BioMart\nclasses: releases/2025-03-16”.

The identities of genes were first converted using the g:Convert tool ^71^ to ensure recognition by the GO schema. The following settings ^49^ were then used for GSEA: minimum set size = 15, maximum set size = 500, minimum intersect number = 2. P-values were corrected for multiple testing across gene sets, based on the gSCS correction method in g:Profiler, separately for each input list of genes corresponding to a given set of subjects (i.e. solved SIT cases, unsolved SIT cases, controls). This method of multiple testing correction takes into account the hierarchical structure of the sets ^71^. We applied a cut-off P-value of adjusted 0.01.

As we had excluded genes from SIT cases when we found a rare variant that passed our filtering criteria in controls, for the control gene list we removed genes with rare variants identified in cases. This reciprocal exclusion produced a control gene list of comparable size to the cases, and therefore a useful negative control for gene set enrichment analysis.

### Functional and structural brain asymmetry

We used the same functional and structural brain asymmetry measures as detailed previously for this dataset, which were available for all stage I and stage II SIT cases and controls ^7,13^.

Briefly, for functional asymmetry, participants had completed four fMRI tasks designed to assess hemispheric lateralization. Two of the tasks typically elicit left-lateralized activation—verbal fluency and tool use pantomiming—while the other two tasks tend to evoke relatively stronger right-hemisphere activation—line bisection judgment and face recognition. Accordingly, the most common pattern is to have two left-lateralized and two right-lateralized brain activation maps ^6^. We used continuous indexes of laterality for each separate task per subject ^7^, as well as a count of the number of atypically lateralized functions per subject (ranging from 0 to 4, where atypical laterality for a task meant a laterality index with opposite sign to the control mean for that task) ^6,7^. Laterality indexes were calculated for each task based on task-appropriate regions of interest and the magnitudes of activation in the top 5% of voxels per hemisphere ^6,7^.

For structural asymmetry, the measures of frontal and occipital petalia and bending were based on an image processing pipeline that involves cortical surface reconstruction, quantifying hemispheric displacement on the anterior-posterior axis, and the angles between the best-fitting plane and the plane x⍰= ⍰0 in the respective lobes ^13,74,75^. Transverse sinus volumes were determined using manual segmentation in ITK-SNAP ^76^, focusing on 20 mm long segments of the left and right transverse sinus that started 5 mm from the midline and ended 25 mm laterally from the midline ^13^.

### Brain imaging genetics

Before sequencing the genomes, it was not known how many or which types of DNA variants would be detected in the dataset, and therefore which group comparisons would be possible to test statistically in relation to brain image metrics. In principle, four main groups were possible: SIT with variants in genes known to cause PCD, SIT with variants in genes not known to cause PCD, SIT with no clear monogenic basis, and controls. In the event, the group of SIT individuals with variants in genes not known to cause PCD was too small for statistical inference (see Results), and all of these genes were anyway cilia-related. We therefore simplified the analysis into three groups: SIT with a likely monogenic basis (genetically ‘solved’), SIT without an obvious monogenic basis (genetically ‘unsolved’), and controls.

#### Functional asymmetries and handedness

To assess group differences in functional brain asymmetries we conducted analyses of covariance (ANCOVA) for each of the four laterality indexes (speech production, manual praxis, spatial attention, and face recognition), where each ANCOVA included a three-level group factor (genetically solved, genetically unsolved, control) as the main predictor, while treating age and sex as covariates:

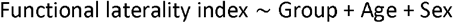

Analyses were performed using the *lm* function in R, and the sum of squares was computed using the *car::Anova* function. Partial eta-squared (*η*^2^) was calculated for the group factor using the ‘effectsize’ package ^77^. To account for multiple comparisons (i.e. four ANCOVAs), Holm-adjusted p-values were calculated.

Furthermore, to assess whether the three groups (solved, unsolved, control) differed in variability of functional asymmetry patterns, we used Levene’s method (median-centered) to test for homogeneity of variance in *N_atypical* (i.e. the total number of atypical functional asymmetry patterns per participant). This was a single test across the three groups.

Additionally, Fisher’s exact test was used to assess whether genetically solved SIT individuals differed in their rate of left-handedness from genetically unsolved SIT individuals. (A comparison was not made with controls, as they were selected to roughly match cases for handedness during study recruitment).

#### Torque measures and transverse sinus volumes

ANCOVA was used to test differences between groups (solved, unsolved, control) in each of four brain torque features (frontal *petalia*, frontal bending, occipital *petalia*, occipital bending), followed by Holm-adjustment of the p-values for the four torque measures tested:

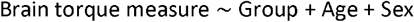

ANCOVA was also used to test for group differences of left and right transverse sinus volumes, with Holm-adjustment for two tests (left and right):

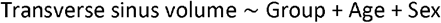

For any ANCOVA tests that were significant with Holm-adjusted p<0.05, we conducted post hoc pairwise comparisons between the solved, unsolved, and control groups. Estimated marginal means were computed using the emmeans package in R, with Tukey’s correction for multiple testing:

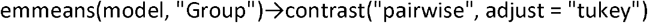

This approach allowed for covariate-adjusted mean comparisons while controlling the familywise error rate. Effect size was estimated with Hedges’ g (bias-corrected standardized mean differences). All analyses were conducted in R (version 4.4.0) using the car (v3.1-3), effectsize (1.0.1), and purr (v1.0.4) packages.

## Results

### Most likely causal DNA variants leading to situs inversus

Among 38 participants with SIT across stages I and II in total, we identified likely causal genetic variants in 19, whom we therefore labelled as genetically ‘solved’ for the purpose of this study (Table 2). Specifically, from stage I, 10 out of 15 SIT participants were genetically solved under a recessive model ^49^, whereas from stage II, 8 out of 23 SIT participants were solved under a recessive model and 1 under a dominant model. Accordingly, 19 cases remained genetically unsolved in total (Table 2).

**Table 2.**
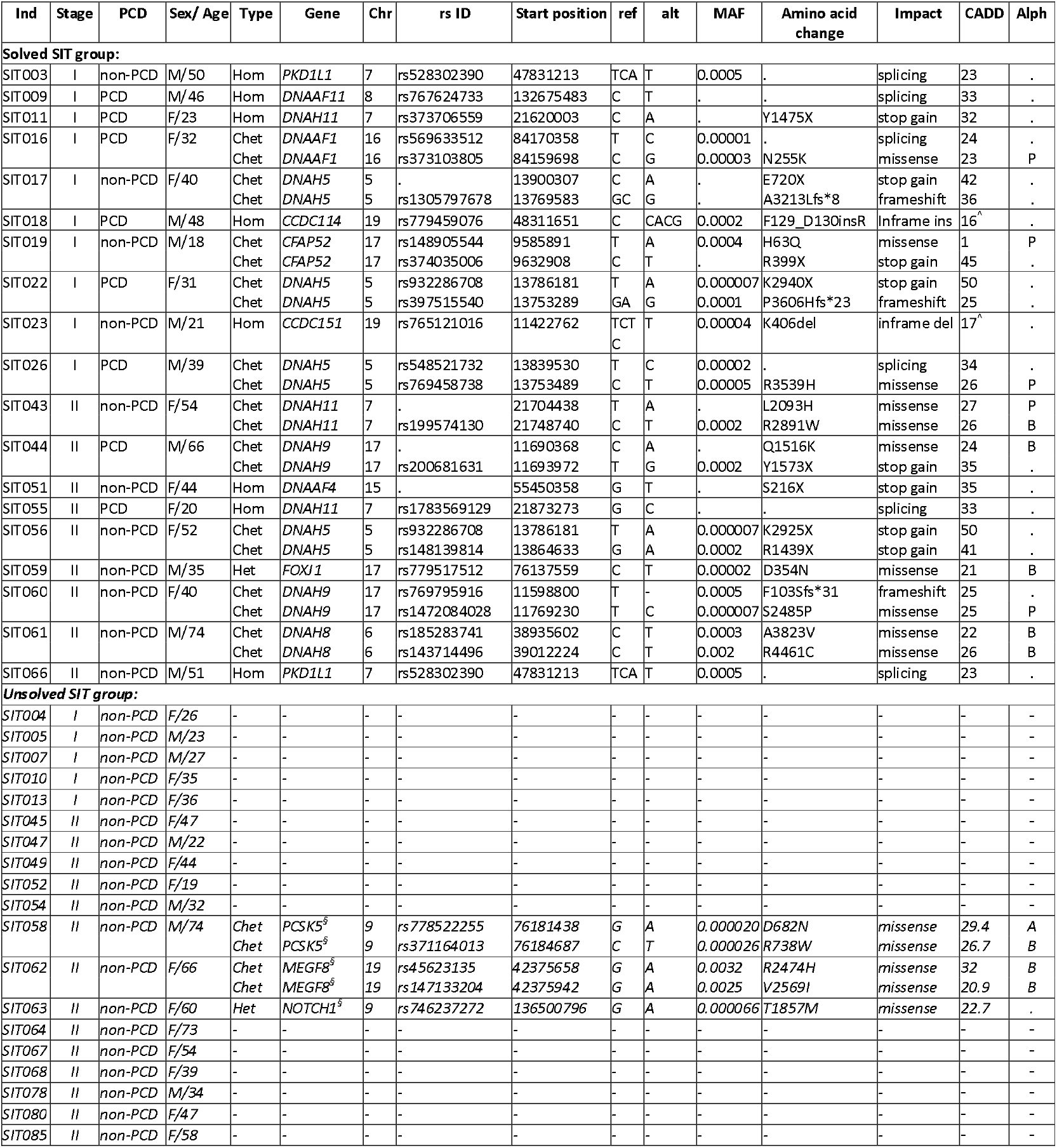
Most likely causal genetic variants for situs inversus in the Belgian dataset. Ind: individual participant. PCD: diagnosis of primary ciliary dyskinesia. M: Male. F: Female. Age: age in years. Hom: homozygous variant, causative under a recessive model. Chet: likely compound heterozygote (causative under a recessive model). Het: heterozygous variant (causative under a dominant model). Chr: chromosome. Ref: allele in the reference genome. Alt: allele present in the study participant. MAF: minor allele frequency (from gnomad4.1_genom_all). Ins: insertion. Del: deletion. CADD (v1.7): combined annotation dependent depletion phred score. Alph: alphamissense annotation. A: Ambiguous. B: Benign. P: Pathogenic. ^§^Stage II finding not conclusive enough to assign causality for human SIT, case considered unsolved (see Results section ‘Notable genetic findings in unsolved SIT individuals’). ^Variant of uncertain clinical significance that was identified as the most likely causal variant by Postema et al. (2019) ^49^ in their earlier analysis of stage I individuals (*CCDC114* and *CCDC151* encode coiled-coil domain containing proteins that are important for the proper function of motile cilia).

Note that the purpose of this study was discovery research on potential links of brain and visceral asymmetries, not clinical diagnosis or feedback to individual participants. Our criteria for assigning likely causal genetic variants were weighted for sensitivity, as appropriate for a phenotype such as SIT which is not, of itself, a pathology and is not necessarily always caused by variants of unambiguous clinical significance. The specificity/sensitivity balance of our criteria could be gaged somewhat by our inclusion of both SIT cases and unaffected controls. In contrast to the 38 SIT cases, of which 19 (50%) were genetically solved according to our criteria for defining causal variants, among the 38 controls only 1 individual met these criteria. Control SIT084 had two missense single nucleotide variants in *DNAH6* with CADD scores >20 (NM_001370.2:exon27:c.G4180A:p.E1394K and NM_001370.2:exon73:c.A11815G:p.S3939G). *DNAH6* encodes a dynein axonemal heavy chain motor protein that drives force and movement within motile cilia and flagella. This finding probably reflects a slight over-sensitivity in our pipeline for the assignment of disruptive status to genetic variants.

The 8 SIT individuals with PCD all had likely disruptive biallelic variants in one gene each that was annotated with ‘Kartagener’, ‘ciliary dyskinesia, primary’ or ‘PCD’ in the ClinVar database ^62^, consistent with a monogenic basis for their phenotype (Table 2). These genes were *DNAAF11* (*LRRC6*) (OMIM: 614930), *DNAH11* (OMIM: 603339), *DNAAF1* (OMIM: 613190), *CCDC114* (OMIM: 615038), and *DNAH5* (OMIM: 603335) from stage I, and *DNAH9* (OMIM: 603330) and *DNAH11* (OMIM: 603339) from stage II. All of these genes encode motile ciliary axoneme components or assembly factors.

There were a further 7 SIT individuals who had no medical record of PCD-like symptoms but did have probable causative biallelic variants for SIT in genes encoding motile ciliary axoneme components or assembly factors, annotated with ‘Kartagener’, ‘ciliary dyskinesia, primary’ or ‘PCD’ in ClinVar (2 SIT individuals from stage I, and 5 from stage II) (Table 2). The implicated genes were *DNAH11* (OMIM: 603339), *DNAAF4* (OMIM: 608706), *DNAH5* (OMIM: 603335), *DNAH9* (OMIM: 603330), and *DNAH8* (OMIM: 603337), and CCDC151 (OMIM: 615956). These findings indicate reduced penetrance for PCD arising from biallelic variants in ciliary genes that can cause SIT ^49,78^.

Three additional non-PCD participants had likely disruptive biallelic variants in genes known to be involved in laterality disorders without PCD. The implicated genes were *CFAP52* (OMIM: 609804) ^79^ and *PKD1L1* (OMIM: 617205) from stage I, and another *PKD1L1* case from stage II (Table 2). CFAP52 protein is part of an axonemal module important for motile ciliary beating ^80^. *PKD1L1* encodes a member of the polycystin cation channel family 1. The potential role of this protein in laterality ^49,81^ may stem from its presence in non-motile cilia of the embryonic node where it mediates a response to leftward fluid flow generated by motile cilia ^82^. However, other roles of PKD1L1 at the node have also been proposed in establishing asymmetry ^83,84^.

One further non-PCD SIT participant had a heterozygous missense variant predicted to be deleterious in *FOXJ1* (OMIM: 602291) (Table 2), encoding a transcription factor and known as an unusual dominant cause of PCD as well as laterality defects ^68,85^.

### Gene set enrichment analysis

For the recessive/X-linked model, the list of genes with disruptive variants according to our criteria, across stages I and II combined, comprised 144 genes in 19 solved SIT individuals, 92 genes in 19 unsolved SIT individuals (Table S2), and 171 genes in 38 controls (Table S2). In solved SIT individuals we expected gene set enrichment analysis to identify ontology sets related to cilia, as each solved individual by definition had such a gene among those that they contributed to the list. Solved SIT individuals would therefore be a useful positive control for the enrichment analysis. For unsolved SIT individuals, gene set enrichment analysis had the potential to discover a biological pathway not previously known to relate to SIT, for example by linking the unsolved SIT individuals through non-ciliary gene sets. The unaffected control individuals provided a negative control for the purpose of gene set enrichment analysis.

As expected, various gene sets related to the ciliary axoneme showed significant enrichment based on genes with recessive causal DNA variants in solved SIT individuals (Table 3). For example, there were sets related to microtubules (microtubule-based movement, microtubule-associated complex) and dynein (dynein complex, dynein intermediate chain binding) (Table 3). In unsolved SIT individuals, no gene sets showed significant enrichment (all gSCS-corrected P>0.01), such that the list of genes with likely causative biallelic DNA variants compiled from these individuals had no discernible commonality (in terms of molecular functions, biological processes or cellular components as defined in the Gene Ontology). In unaffected controls, as expected, no gene sets showed significant enrichment (all gSCS-corrected P>0.01).

**Table 3.** Results from gene set enrichment analysis. GO: gene ontology. ^*^gSCS corrected p values (see Methods).

| Group | GO term name | Enrichment p_value* | Term size | Query size | Overlap size | Intersection |
| --- | --- | --- | --- | --- | --- | --- |
| SIT solved | cytoplasmic region | 3.8x10<sup>-6</sup> | 313 | 118 | 13 | CLASP2, DCTN1, DNAAF1, DNAH11, DNAH12, DNAH5, DNAH8, DNAH9, DST, HIF1A, LRRK2, PCLO, RP1L1 |
|  | minus-end-directed microtubule motor activity | 1.3x10<sup>-5</sup> | 17 | 107 | 5 | DNAH11, DNAH12, DNAH5, DNAH8, DNAH9 |
|  | microtubule-based movement | 3.4x10<sup>-5</sup> | 478 | 106 | 15 | CELSR2, DNAAF1, DNAAF4, DNAH11, DNAH12, DNAH5, DNAH8, DNAH9, DST, FMN2, HIF1A, KIF13B, NEFH, STARD9, SYNE2 |
|  | microtubule motor activity | 4.2x10<sup>-5</sup> | 68 | 107 | 7 | DNAH11, DNAH12, DNAH5, DNAH8, DNAH9, KIF13B, STARD9 |
|  | cytoskeletal motor activity | 1.0x10<sup>-4</sup> | 115 | 107 | 8 | DNAH11, DNAH12, DNAH5, DNAH8, DNAH9, KIF13B, MYH1, STARD9 |
|  | dynein complex | 1.3x10<sup>-4</sup> | 54 | 118 | 6 | DCTN1, DNAH11, DNAH12, DNAH5, DNAH8, DNAH9 |
|  | polypeptide conformation or assembly isomerase activity | 1.5x10<sup>-4</sup> | 121 | 107 | 8 | DNAH11, DNAH12, DNAH5, DNAH8, DNAH9, KIF13B, MYH1, STARD9 |
|  | dynein light intermediate chain binding | 2.0x10<sup>-4</sup> | 28 | 107 | 5 | DNAH11, DNAH12, DNAH5, DNAH8, DNAH9 |
|  | plasma membrane bounded cell projection cytoplasm | 7.0x10<sup>-4</sup> | 277 | 118 | 10 | DNAAF1, DNAH11, DNAH12, DNAH5, DNAH8, DNAH9, DST, HIF1A, LRRK2, RP1L1 |
|  | dynein intermediate chain binding | 8.6x10<sup>-4</sup> | 37 | 107 | 5 | DNAH11, DNAH12, DNAH5, DNAH8, DNAH9 |
|  | axonemal dynein complex | 0.0015 | 22 | 118 | 4 | DNAH12, DNAH5, DNAH8, DNAH9 |
|  | microtubule plus-end | 0.0031 | 26 | 118 | 4 | CLASP2, DCTN1, DST, NCKAP5 |
|  | microtubule-based transport | 0.0033 | 220 | 106 | 9 | DNAAF1, DNAAF4, DNAH11, DNAH5, DNAH9, DST, HIF1A, NEFH, SYNE2 |
|  | microtubule | 0.0038 | 492 | 118 | 12 | CLASP2, DCTN1, DNAH11, DNAH12, DNAH5, DNAH8, DNAH9, DST, KIF13B, NCKAP5, RP1L1, SHROOM2 |
|  | epithelial cilium movement involved in extracellular fluid movement | 0.0052 | 44 | 106 | 5 | DNAAF1, DNAAF4, DNAH11, DNAH5, DNAH9 |
|  | microtubule associated complex | 0.0069 | 159 | 118 | 7 | DCTN1, DNAH11, DNAH12, DNAH5, DNAH8, DNAH9, KIF13B |
|  | extracellular transport | 0.0074 | 47 | 106 | 5 | DNAAF1, DNAAF4, DNAH11, DNAH5, DNAH9 |
| SIT unsolved | No significant enrichment (all gSCS corrected p values > 0.01) |  |  |  |  |  |
| Control | No significant enrichment (all gSCS corrected p values > 0.01) |  |  |  |  |  |

For the dominant model, there were 860 genes with potential causal variants according to our criteria in unsolved SIT individuals, combined across stages I and II (Table S2). This number of genes was too high relative to the number of unsolved individuals (19) to perform gene set enrichment analysis with respect to a monogenic model.

### Notable genetic findings in unsolved SIT individuals

In this section we mention notable genetic findings in unsolved SIT individuals without PCD, some of which may putatively have led to their condition. We did not consider these findings conclusive enough to assign causality, but this information may be valuable in combination with future studies of SIT.

In SIT individual SIT058, we found potential compound heterozygous missense variants in *PCSK5* (OMIM: 600488) (see Table 2 for information on the exact variants). This gene encodes a proprotein convertase enzyme. In mice, *PCSK5* disruption is linked to heterotaxy, dextrocardia and abnormal respiratory motile cilium physiology ^86^, as well as the ‘VACTERL association’ (a sporadic congenital disorder that includes vertebral defects, anal atresia, cardiac malformations, esophageal atresia, radial or renal dysplasia and limb anomalies) ^87^. Human *PCSK5* has also been linked to the VACTERL association (OMIM: 192350) ^87^, but unsolved SIT individual SIT058 did not present with a cluster of phenotypes resembling VACTERL. To our knowledge, ours is the first study to find a putative effect of *PCSK5* in a human SIT individual, and we are therefore cautious about assigning causality to this finding.

In unsolved SIT individual SIT062, potential compound heterozygous missense variants in *MEGF8* (OMIM: 604267) were identified (Table 2). This gene encodes a transmembrane protein involved in negative regulation of Hedgehog signaling, a pathway involved in embryonic cell differentiation ^88^. In mice, biallelic mutations in *MEGF8* can cause heterotaxy and complex congenital heart defects, whereby Nodal signaling is disrupted during embryonic left-right axis formation, but with unaffected nodal cilia motility ^89^. In humans, biallelic pathogenic variants in *MEGF8* cause Carpenter syndrome-2 (CRPT2; OMIM 614976), a multiple congenital malformation disorder characterized by multisuture craniosynostosis and polysyndactyly of the hands and feet, often accompanied by abnormal left-right patterning including situs inversus, and other features such as obesity, umbilical hernia, cryptorchidism, and congenital heart disease ^90,91^. As SIT individual SIT062 had no indication of the Carpenter syndrome-2 multiple phenotype beyond SIT, we cannot confidently assign *MEGF8* as causal to SIT in this individual. However, the *MEGF8* missense variants that we identified may plausibly have contributed to SIT in this individual, with reduced penetrance for a broader phenotype.

In unsolved SIT individual SIT063, we found a rare heterozygous missense variant predicted to be deleterious in *NOTCH1* (OMIM: 190198). The Notch signaling pathway is involved in processes related to cell fate specification, differentiation, proliferation, and survival, and Notch activity helps to establish left-right asymmetry in chick and mouse embryos through affecting asymmetric gene expression around the node ^92,93^. In humans, pathogenic *NOTCH1* variants are associated with autosomal dominant non-syndromic congenital heart disease and sometimes Adams–Oliver syndrome 5 (OMIM: 616028), a rare developmental disorder defined by scalp and limb defects and often accompanied by vascular and heart defects ^94^. However, these *NOTCH1*-associated conditions in humans have not been characterised as laterality disorders to our knowledge, and we cannot therefore assign causality for the SIT of individual SIT063.

For all the gene sets that showed significant enrichment from genetically solved SIT cases, we cross-referenced their member genes with the genes carrying disruptive variants according to our criteria in unsolved cases. There were 12 genes showing overlap: *CCDC66, CFAP77, CLIP2, DCTN1, DNAI7, EML6, KATNAL2, MAP2, MYH7B, MYO7B, RIMS2*, and *RP1L1*. On further literature and database searching, none of these genes were clear candidates for causing human SIT (for example, human pathogenic variants were already known to cause other phenotypes not including disrupted laterality, or we only observed heterozygous variants in genes previously linked to recessive phenotypes).

The maximum number of unsolved SIT cases with disruptive variants in the same gene was 3, concerning potential deafness-related genes *NCOA3* (OMIM 601937) (individuals SIT052, SIT067 and SIT085) and *UMODL1* (OMIM 613859) (individuals SIT052, SIT080 and SIT085). Neither of these genes has any prior reported link to disrupted visceral laterality.

### Functional brain asymmetry

In ANCOVAs across the genetically solved, unsolved and control groups, no significant effects of group were found on the functional laterality indexes for word generation (*F*_*(2, 71)*_ = 2.75, *p*_*adjusted*_ = 0.21, *η*^2^ = 0.07), spatial attention (*F_(2, 71)_* = 4.35, *p*_*adjusted*_ = 0.07, *η*^2^ = 0.11), praxis (*F_(2, 71)_* = 1.13, *p*_*adjusted*_ = 0.44, η^2^ = 0.03), or face recognition tasks (*F*_*(2, 71)*_ = 1.55, *p*_*adjusted*_ = 0.44, *η*^2^ = 0.04). The distributions of functional brain asymmetry indexes in the different groups are shown in Figure 1.

**Figure 1.**
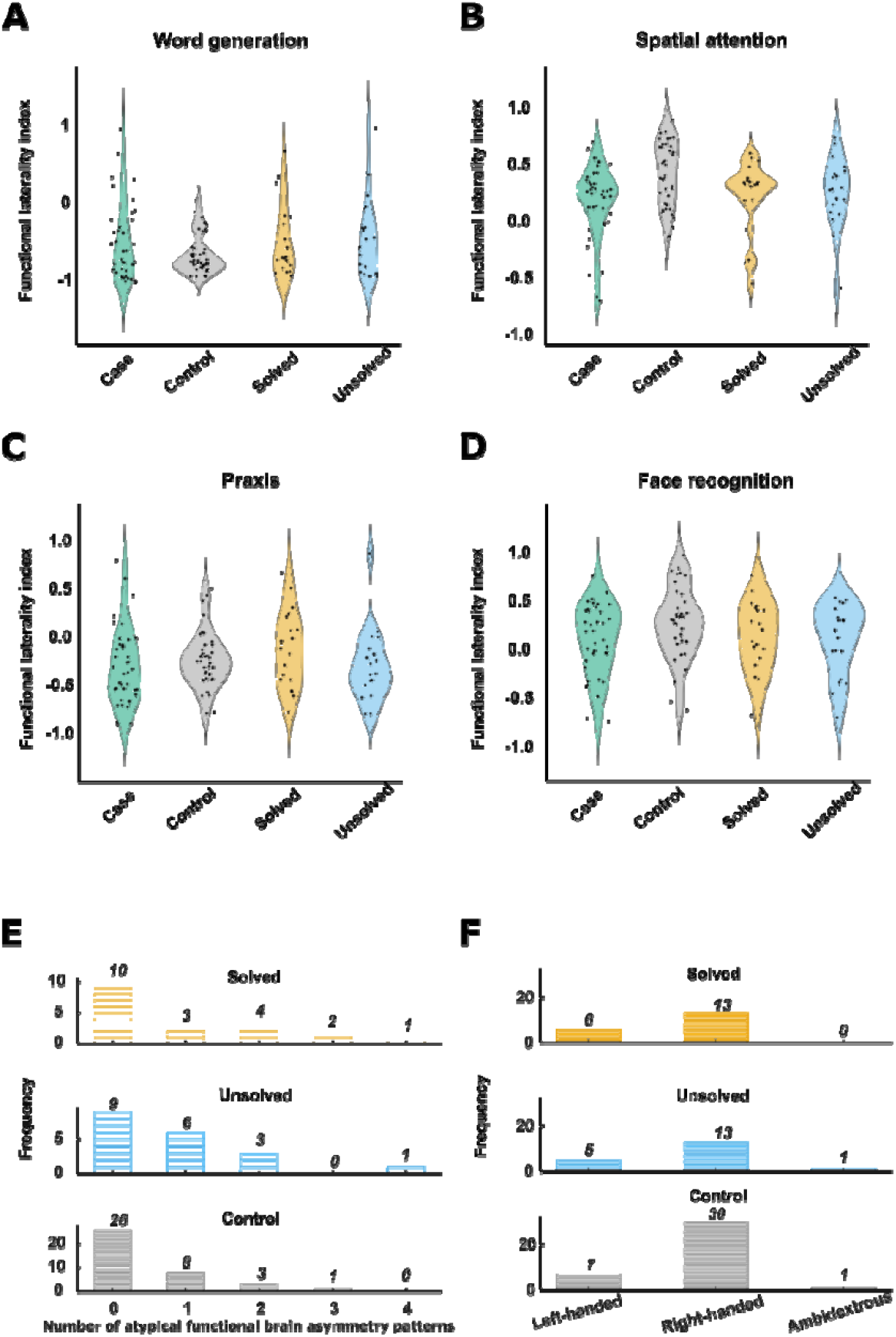
Functional brain asymmetry and handedness in 38 SIT individuals and 38 controls. **A-D**. Functional brain laterality indexes for four different tasks. Data from SIT individuals are shown as one group all together, and also stratified into genetically solved and unsolved groups. Positive values on the y-axis represent rightward functional lateralization while negative values represent leftward functional lateralization. **E**. The number of atypically lateralized functions among genetically solved, unsolved, and control groups. **F**. Handedness in the solved, unsolved, and control groups. (During study recruitment, controls were roughly matched for handedness with SIT cases, therefore we only tested for the group difference between solved and unsolved SIT cases.)

For the number of atypically lateralized functions (ranging from 0 to 4), there was no significant difference in variance across the genetically solved, unsolved and control groups (Levene’s test, *p* =.085) (Figure 1). The rate of left-handedness did not differ between genetically solved and unsolved SIT groups (Fisher’s exact *p* = 1) (Figure 1).

For genes that were recurrently implicated in more than one solved SIT individual, there was no discernible pattern in terms of typical or atypical functional brain asymmetry. For example, among four SIT individuals with biallelic causal variants in *DNAH5* (a dynein heavy chain gene involved in ciliary motion), two individuals showed typical functional asymmetry for all tasks, while the other two showed 2 or 3 atypically lateralized functions (Table S3). Both of the SIT individuals with biallelic *DNAH9* causal variants showed typical functional laterality for all four tasks. Of the three SIT individuals with biallelic *DNAH11* causal variants, two showed no atypically lateralized functions and one showed 1 atypically lateralized function. Of the two SIT individuals with biallelic *PKD1L1* causal variants (a known cause of SIT without PCD), one individual showed no atypically lateralized functions and one showed 2 atypically lateralized functions.

Furthermore, of the two individuals with atypical functional brain asymmetry for all four tasks, one was genetically unsolved, and the other had biallelic causal variants in *CCDC151* (which encodes an axonemal protein required for attachment of outer dynein arm complexes to microtubules in motile cilia).

In summary, we could not link causal heterogeneity of SIT to functional brain asymmetry in this dataset.

### Structural brain asymmetry

ANCOVAs revealed significant associations between the three-level group factor (genetically solved SIT cases, unsolved SIT cases, and controls) and frontal petalia (*F*_*(2, 71)*_ = 4.63, *p*_*ajusted*_ =.03, *η*^2^_partial_= 0.12), occipital petalia (*F*_*(2, 71)*_ = 8.13, *p*_*ajusted*_ =.002, *η*^2^_partial_ = 0.19), and occipital bending (*F*_*(2, 71)*_ = 10.6, *p*_*ajusted*_ =.0003, *η*^2^_partial_ = 0.23) (Figure 2), indicating robust differences across groups in these brain torque features. The ANCOVA for frontal bending was not significant (*F*_*(2, 71)*_ = 1.01, *p* =.37, *η*^2^_partial_ = 0.025) (Figure 2).

**Figure 2.**
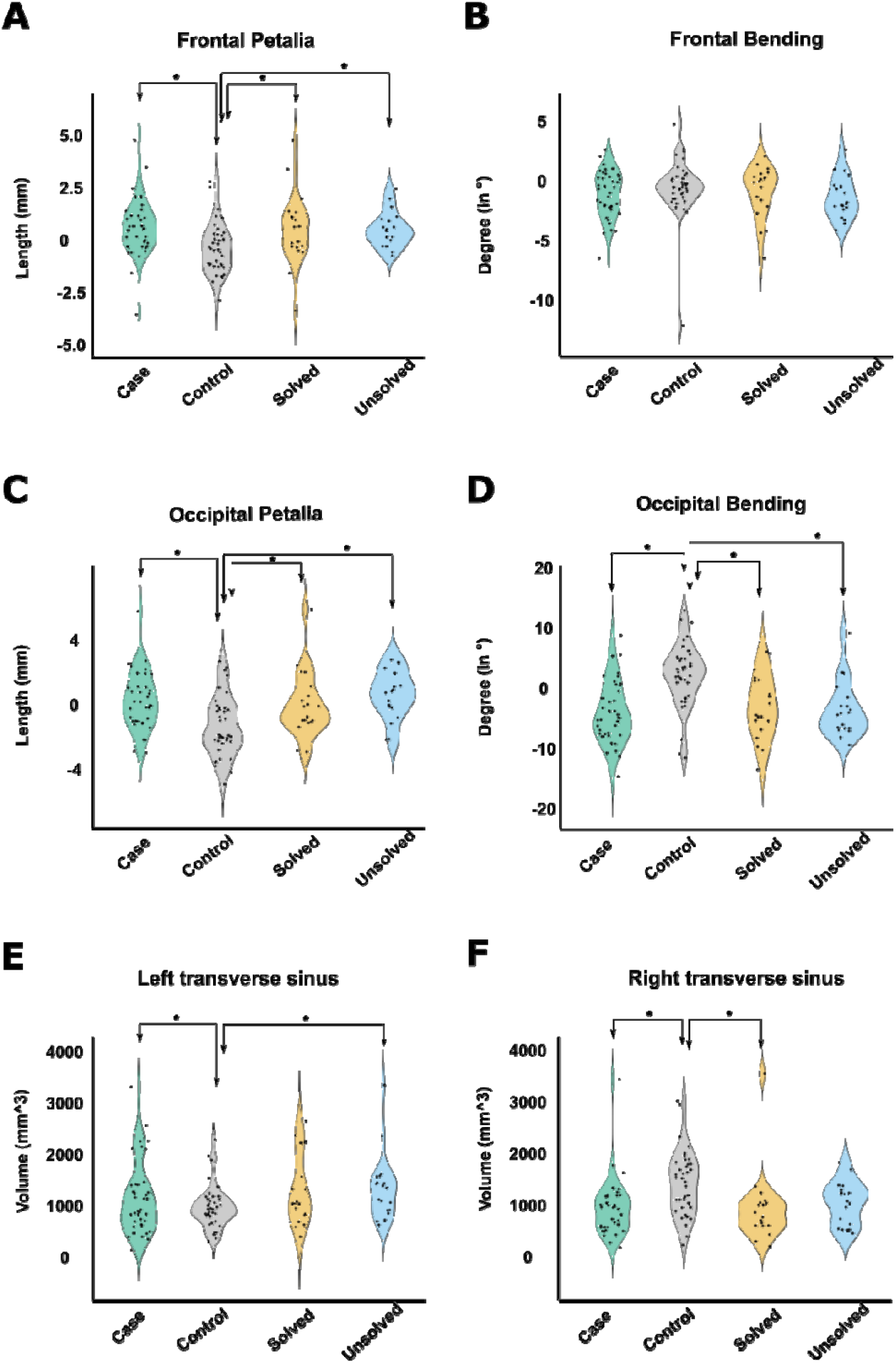
Brain torque metrics and transverse sinus volumes in 38 SIT cases and 38 controls. **A-F** Distributions of structural measures in different groups. Data from SIT individuals are shown as one group all together (labelled ‘Case’) and also stratified into genetically solved (n=19) and unsolved (n=19) groups. ^*^Indicates a nominally significant group difference (p<0.05). While various differences were found between SIT cases and controls, no differences were found between solved and unsolved SIT cases.

However, post hoc pairwise comparisons revealed that significant effects in ANCOVA were driven by differences between solved SIT cases and controls (frontal petalia, effect size = 0.70, *p* =.04; occipital petalia, effect size = 0.74, *p* =.03; occipital bending, effect size = 1.01, *p* =.002) and also differences between unsolved SIT cases and controls (frontal petalia, effect size = 0.70, *p* =.04; occipital petalia, effect size = 1.06, *p* =.001; occipital bending, effect size = 1.10, *p* =.002), but not by differences between solved SIT cases and unsolved SIT cases (frontal petalia, effect size = 0.003, *p* = 1.00; occipital petalia, effect size = 0.32, *p* =.61; occipital bending, effect size = 0.10, *p* =.95) (Figure 2). In other words, brain torque features differed on average between SIT cases and controls regardless of the genetic status of cases, while torque features did not differ between genetically solved and unsolved SIT cases.

ANCOVAs revealed significant associations between the three-level group factor (solved SIT, unsolved SIT, control) and both left transverse sinus volume (*F*_*(2, 71)*_ = 3.97, *p*_*ajusted*_ =.02, *η*^2^_partial_ = 0.10) (Figure 2) and right transverse sinus volume (*F*_*(2, 71)*_ = 4.96, *p*_*ajusted*_ =.02, *η*^2^_partial_ = 0.12) (Figure 2), indicating volumetric differences across the groups. However, again post hoc comparisons revealed no significant differences between the solved and unsolved SIT cases (left transverse sinus, effect size = 0.14, *p* =.91; right transverse sinus, effect size = 0.18, *p* = 0.86), such that the significant ANCOVAs were again driven by differences between SIT cases and controls, and not by causal heterogeneity between cases (Figure 2).

## Discussion

SIT is a rare developmental syndrome that is monogenic in some affected individuals. In this study we analyzed the largest brain imaging genetic dataset of SIT ever assembled. Through genome sequencing to identify the most likely causative DNA variants in each participant, we solved 19 SIT cases with predominantly biallelic disruptive variants in ciliary genes, while 19 SIT cases remained genetically unsolved. This study had the potential to link the causal heterogeneity of SIT to variations in brain functional and structural asymmetry. Instead, we discovered that regardless of the presence or absence of an overt genetic lesion affecting ciliary proteins, SIT is not obviously associated with functional brain asymmetry, but is associated with altered structural brain torque. These findings shed new light on the genetics and development of brain asymmetry in humans.

In terms of functional asymmetry across four cognitive domains, genetically solved and unsolved SIT cases did not differ on average from each other, nor from controls. This means that typical population asymmetries such as left-hemisphere language dominance are mostly unaltered in SIT, both when SIT is caused by overt monogenic disruptions of ciliary function and in their absence. Therefore, altered embryonic left-right axis formation that patterns visceral organ asymmetry, regardless of its cause, appears to have little effect on functional brain asymmetry. The unknown mechanisms for patterning functional brain asymmetry in the human brain appear largely independent of the ciliary-nodal pathway, but also robust to potential environmental or random developmental perturbations of embryonic visceral situs, to the extent that such factors lead to SIT in the absence of overt monogenic causes. Interestingly, this contrasts to the situation in zebrafish, where visceral situs inversus and the nodal pathway are linked to both neuroanatomic and behavioural reversals on the left-right axis ^95^.

As left-hemisphere dominance for language is present in roughly 85% of people, our findings suggest a strong developmentally lateralized bias in functional brain asymmetry that is distinct from the ciliary-nodal pathway and also largely unaffected by non-genetic risk factors for SIT. Consistent with this, we have proposed an organ-intrinsic mechanism of brain asymmetry based on cellular chirality and organized by microtubules, but distinct from those affecting motile ciliary function at the node ^31,33,34^. This would be compatible with the implication of microtubules in brain and behavioural asymmetries by large-scale genome-wide association studies ^31-36^, together with the fact that functional brain asymmetry is mostly unaltered in people with SIT. The present study adds to this understanding, by showing that altered functional brain asymmetry is not a group-level feature even when SIT is broken down by the presence or absence of identifiable causal genetic variants.

It is important to note that PCD-causing DNA variants do not always result in SIT, but instead lead to randomization of left-right axis formation. This means that roughly half of carriers of biallelic variants affecting genes such as *DNAH5* or *DNAAF11* (encoding motile ciliary axoneme components or assembly factors) have SIT, while the other half have unaltered visceral situs ^37^. In principle, randomization might also play a role in the development of functional brain asymmetry ^42,43,96^, whenever mechanisms of establishing typical brain asymmetry are disrupted. Hence, we not only tested for group mean effects on functional brain asymmetry, but also tested whether the variance in the number of atypically lateralized functions might differ between genetically solved SIT, unsolved SIT and control individuals. We did not find differences in variance, again underscoring a large degree of independence between functional brain asymmetry and visceral situs placement. This independence is now known from the present study to occur regardless of the precise mechanism of visceral situs perturbation.

For example, it may be that in genetically unsolved SIT cases, developmental perturbations of visceral situs placement occur further downstream of nodal ciliary action, perhaps directly affecting gene expression asymmetries of key signaling molecules or transcription factors in the visceral left-right differentiation pathway. If left-side and right-side concentrations of key signaling molecules do not differ sufficiently at key moments, and/or in key cell types, during the establishment of visceral organ situs in the early embryo, then situs might be established closer to the chance 50:50 level. The same principle could apply to functional brain asymmetry even when mechanisms are largely distinct from visceral situs.

However, we cannot rule out that certain rare genetic lesions might impact both visceral and functional brain asymmetries, as no more than four individuals in our dataset had likely causative DNA variants in the same gene (Table 2). This number is too small to associate altered functional brain asymmetry with genetic factors when grouped by individual genes, especially when randomization may play a role within such a group. We note also that left-handedness – another aspect of functional brain asymmetry – is somewhat over-represented among the 38 SIT individuals studied here, compared to the general population ^7^ (but not compared to the controls in the present study who were selected for handedness roughly balanced with the SIT cases). Furthermore, the gene *NME7*, which encodes a γ-tubulin-associated protein, has been implicated in SIT ^97^, handedness and surface area asymmetry of the anterior insular cortex ^34^. The present study did not identify a SIT individual with this specific gene affected, but *NME7* provides a potential mechanistic link between visceral and brain asymmetries.

Furthermore, regarding structural brain torque and transverse sinus volumes, the evidence clearly suggests a developmental link with visceral situs. Most strikingly, the direction of occipital bending is reversed in SIT at the group level compared to controls in the Belgian dataset (Figure 2). The present study revealed that this group-level alteration of brain torque features is found in SIT regardless of the presence or absence of likely causative DNA variants in ciliary genes. Thus, we envision a fairly tight developmental coupling of brain torque with visceral situs through shared developmental mechanisms, possibly mediated via the anatomy of the posterior venous system ^13^. It also follows that functional brain asymmetry for e.g. hemispheric language dominance is largely decoupled from brain torque. This is also consistent with the association between brain torque and handedness in the general population having only a small effect size ^17^.

Eight of our 38 SIT cases had medical records indicating PCD, which involves impaired motile ciliary function. All eight of these cases were solved by biallelic variants disrupting genes known to cause PCD and SIT. However, an additional 8 SIT cases with no medical records indicating PCD were found to have likely causative variants in PCD-linked genes that encode motile ciliary axoneme components or assembly factors (*DNAH5, DNAH8, DNAH9, DNAH11, DNAAF4, CCDC151*) or the PCD-linked transcription factor *FOXJ1*. Therefore, causative variants in these genes appear to have reduced penetrance for PCD ^49,78^, and a diagnosis of PCD may not be a good indicator for the presence/absence of disrupted nodal motile ciliary function. Genome sequencing was therefore important for the present study, to properly assess the genetic causation of SIT in relation to brain asymmetry.

Through gene set enrichment analysis of genes with disruptive biallelic variants in unsolved SIT cases, this study had the potential to implicate a novel pathway in the etiology of SIT, and perhaps then to link a newly defined subgroup of SIT to the presence/absence of altered brain asymmetry. In the event, unsolved SIT cases showed no discernible commonality in terms of pathways affected by the gene variants that they carried. This makes non-genetic causes more plausible explanations for the unsolved cases, such as chance and/or exposure *in utero* to SIT risk factors such as maternal smoking. It remains possible that our variant calling and filtering pipeline was not sufficiently sensitive to detect all causative genetic variants in the Belgian SIT sample, but tuning these parameters to improve sensitivity would likely also cause a drop in specificity (we already observed one control individual having biallelic *DNAH6* disruptive variants as defined under our current pipeline, which we suspect to have little impact on the protein in this individual).

Our focus on exonic variants (substitutions and insertion-deletions) may have missed other types of monogenic causes for SIT in unsolved cases, such as larger genomic copy number variants (CNVs) or variants outside coding regions. Oligogenic or polygenic effects may also be involved. Searching for CNVs in unsolved SIT cases is tractable and may be fruitful in future studies, whereas much larger sample sizes would likely be needed to screen through the non-coding genome or address multigenic causation. As SIT only occurs in roughly 1 out of every 10,000 people and is not necessarily a pathogenic trait unless accompanied by another condition, then a large-scale genetic study does not appear feasible until/unless genome sequencing becomes a standard part of ordinary healthcare.

Future studies would benefit from considering more aspects of brain asymmetry than assessed here, including structural brain asymmetry beyond torque (such as cerebral cortical regional surface area and thickness asymmetries and subcortical volume asymmetries), as well as asymmetries of structural and functional connectivity in the brain. Previous studies in smaller subsets of the Belgian SIT dataset have found no clear group differences between SIT individuals and controls in terms of white matter asymmetries ^98^ or structural asymmetries in and around the Sylvian fissure ^99^. However, analysis of the total combined dataset has not been performed for these brain metrics, nor has genetic data been considered in relation to them.

In conclusion, the present study indicates that altered functional brain asymmetry is not generally a feature of SIT, regardless of whether or not SIT cases carry disruptive variants in ciliary genes. This indicates a large degree of developmental independence between functional brain asymmetry and visceral organ asymmetry, and leaves the field still searching for mechanisms that might pattern the brain to be functionally asymmetric for e.g. language in most people. This study also found that cerebral torque (an aspect of asymmetrical brain structure) is affected in people with SIT, but again regardless of whether or not causative variants in ciliary genes are present. Therefore, brain torque is closely developmentally tied to visceral asymmetry, to the extent that genetic or non-genetic perturbations may both impact a shared pathway that affects body and brain structure. It also follows that functional brain asymmetries develop largely independently of brain anatomical torque.

## Acknowledgements

This research was funded by the Max Planck Society (Germany) and the Fonds Wetenschappelijk Onderzoek Vlaanderen (FWO-grant n⍰ G.0114.16 N assigned to Guy Vingerhoets).

## Data availability

The genome sequence data cannot be freely available due to privacy/identifiability concerns and the fact that consent was not given for such dissemination. Please contact the corresponding author regarding data access. The brain imaging data are not publicly available due to privacy regulations but are available upon reasonable request. The demographic and image-derived metrics analyzed in the present study are available for download from https://osf.io/w48z2/?view_only=4d527d7bac13436b9989ecb85ebeebb6 and https://link.springer.com/article/10.1007/s00429-017-1598-5. Code used in the present study can be accessed here: https://github.com/MengYunWang/Situs_inversus

## Author contributions

Conceptualization: M-YW, SEF, GV, CF. Methodology: M-YW, NN, EE. CF. Software: M-YW. Formal analysis: M-YW. Investigation: M-YW, CF. Resources: SEF, GV, CF. Data curation: M-YW. Visualization: M-YW. Supervision: CF. Project administration: CF. Funding acquisition: SEF, GV, CF. Writing - original draft: M-YW, CF. Writing - review and editing: all authors.

